# bayesTPC: Bayesian inference for Thermal Performance Curves in R

**DOI:** 10.1101/2024.04.25.591212

**Authors:** Sean Sorek, John W. Smith, Paul J. Huxley, Leah R. Johnson

## Abstract

1. Reliable predictions of arthropod responses to climatic warming are important because many of these species perform important roles that can directly impact human society.
2. Thermal performance curves (TPCs) provide useful information on the physiological constraints that limit the capacity of temperature-sensitive organisms (like arthropods) to exist and grow.
3. NLS pipelines for fitting TPCs are widely available, but these approaches rely on assumptions that can yield unreliable parameter estimates.
4. We present `bayesTPC`, an `R` package for fitting TPCs to trait responses using the nimble language and machinery as the underlying engine for Markov Chain Monte Carlo. `bayesTPC` aims to support the adoption of Bayesian approaches in thermal physiology, and promote TPC fitting that adequately quantifies uncertainty.

## 2 Introduction

Predicting arthropod responses to climatic warming is important because these species perform critical roles as pollinators, disease vectors, and agricultural pests. So far, research effort has focused on determining species’ lethal temperatures (high or low temperature limits) [e.g., 1, 2] and predicting specific temperatures at which their thermal fitness is optimized [e.g., 3]. Thermal performance experiments measure traits (e.g., juvenile development time) across a range of constant temperatures to obtain thermal performance curves (TPC) [e.g., 4, 5]. Specific mathematical functions are often used as TPCs to quantify the full shape of the thermal response [6]. Many possible shapes (e.g., symmetric, left/right-skewed) are available through different proposed TPC functions [see 7, 8].

TPCs are typically fitted to trait data using non-linear least squares [NLS, e.g., rTPC, 9]. This choice is pragmatic – NLS pipelines are widely available, and, if optimization routines work, fitting occurs quickly. However, this approach has several drawbacks. First is that NLS algorithms require good initial guesses. Multi-start schemes [e.g., 9] can ameliorate this issue, but they cannot guarantee convergence to a global optima. Moreover, NLS assumes that observed data are normally distributed with constant variance. This assumption suits many values, but it can be unreasonable for data with well-defined bounds, high skew, or non-constant variance. Such assumption mismatch can yield poorly estimated parameters and biased predictions. Further, NLS fits typically use normal approximations for the uncertainty in parameters/confidence intervals and for predictive intervals. Normal distributions have infinite support, so biologically unrealistic predictions (such as negative traits or parameter bounds) can be made. Bootstrapping can solve this issue [e.g., in `rTPC`, 9], but tends to be unstable for small datasets.

Likelihood-based methods, including Bayesian approaches, can ameliorate many of the issues with NLS. For example, formal definitions of data distributions reduces the potential for NLS assumptions to yield unreliable parameter estimates. They allow model comparison through information theory [e.g., 10]. Bayesian approaches additionally allow straightforward, principled inclusion of biological knowledge into analyses while providing multiple uncertainty estimates, even for small datasets.

Bayesian approaches have previously been used to fit TPCs for vector-borne disease systems [e.g., 11, 12]. Although training and code for researchers to employ Bayesian approaches are available (e.g., VectorBiTE RCN), their adoption remains slow because the barrier to entry remains high. Accordingly, we developed the `bayesTPC` package to allow simple Bayesian TPC fitting in `R`. We include tools to assist output inspection, provide summaries, and plot results. Our package release includes implementations of multiple TPCs and simple Bayesian generalized linear models (GLMs), including sensible default priors for TPCs suitable for small arthropods.

Below we introduce the general approach to Bayesian TPC fitting and its implementation in `bayesTPC`. We outline package functionality, and provide a simple example of how the package can be used. Finally, we discuss some of our specific choices and subtleties for users to bear in mind.

## 3 Conceptual Approach

`bayesTPC` attempts to estimate the probability distribution of the parameters, ***θ***, of a specific TPC and observation model given a data set of a performance trait **Y**_obs_ across a range of temperatures `T` while accounting for data uncertainty. We assume that the observed data represent underlying traits that can be modeled as a function of the parameters and temperatures, ***f***, together with an observation model ℳ:

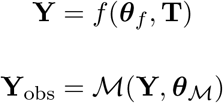

where *θ*_*f*_ are the TPC specific parameters, and *θ* _*ℳ*_ are the parameters of the observation model (so that ***θ*** = (*θ*_*f*_, *θ*_*ℳ*_). For example, if there is additive Gaussian noise with mean *θ* and variance *σ*^2^, then the model is:

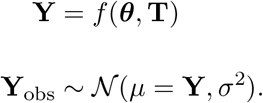

Our inference target is the posterior distribution of parameters (or simply the posterior distribution), Pr(*θ*|*𝒴*), where Pr(*·*) denotes a probability, 𝒴 denotes the data (both traits and temperatures) and *θ* denotes the set of model parameters (both for the TPC and the observation model). Using Bayes theorem, the posterior distribution is calculated as:

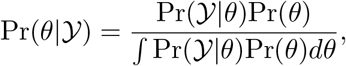

where Pr(𝒴|*θ*) is the likelihood and Pr(*θ*) is the prior, which encodes the prior knowledge about the parameters before data collection. The denominator is the probability of observing the data under any settings of model parameters. Given observed data, the denominator is simply a scaling constant.

For sufficiently complex models, a closed form solution of the posterior distribution is not obtainable. Instead, the posterior distribution is approximated using methods like Markov Chain Monte Carlo (MCMC) [13]. MCMC approximates the posterior by taking pseudo-random samples from the target distribution. With these we can obtain the posterior of any function of the parameters [14] through the “plug-in principle”. For example, the TPC posterior distribution can be obtained, as can the posterior predictive distribution (including both uncertainty due to parameter estimates and the noise model).

Many flexible, general use tools facilitate Bayesian model building and fitting, reducing the necessity to code MCMC samplers from scratch [e.g., JAGS, 15; `nimble`, 16]. Using such tools is easier than implementing MCMC directly, but they are inaccessible for inexperienced users. The fact that NLS is still more common than Bayesian TPC fitting (via JAGS) despite them being been available since 2015 [11] reflects this. Indeed, new NLS pipelines for TPC fitting continue to be published [9], but Bayesian pipelines remain rare.

## 4 Technical Overview

### 4.1 Installation

`bayesTPC` can be installed using `remotes::install_github(“johnwilliamsmithjr/bayesTPC”)`. Note that `nimble` [16] must be installed and loaded before `bayesTPC` can be used.

### 4.2 Thermal Performance Curve (TPC) Models

`bayesTPC` uses a proprietary model specification format which stores the model formula, likelihood, and priors. We include eight common TPC models (listed with `get_models()`) in the package (Table 1). Users can obtain the default specification and priors for these models with `get_default_model_specification()`. The package also includes linear and quadratic formulations for Bernoulli, binomial, and Poisson GLMs, assuming canonical link functions.

**Table 1.**
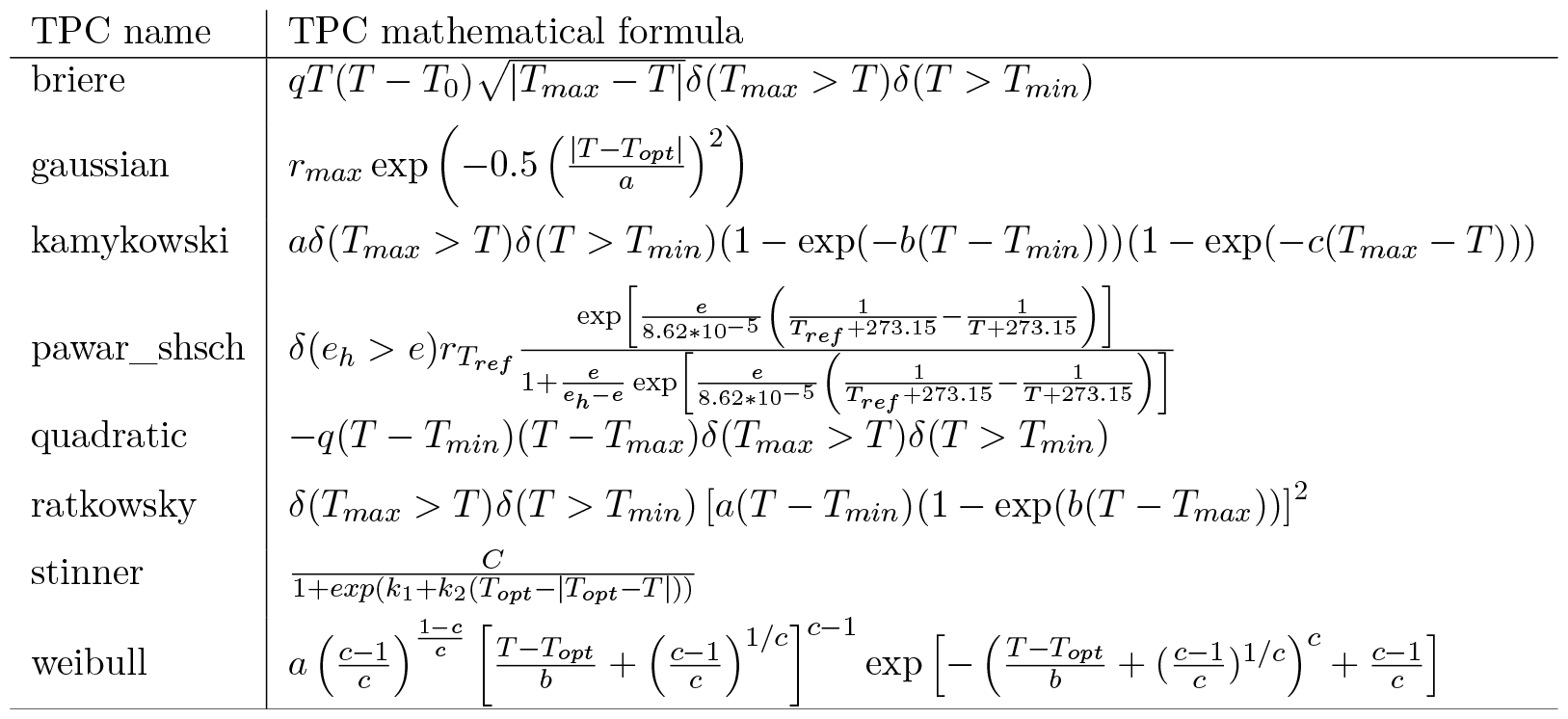
Thermal performance curves included in bayesTPC.

### 4.3 Fitting

TPC models are fitted using the `b_TPC()` function, which requires two user-specified inputs. The first is `data`; a list with expected entries named “Trait” corresponding to the trait being modeled by the TPC and “Temp” corresponding to the temperature in Celsius that the trait was measured at. The second input is `model`, which is a string specifying the model name or a `btpc_model` object. If a string is provided, the default model specification is used.

The default specification for each model includes a set of default priors for all TPC and observation parameters, including a default likelihood (a truncated normal distribution). Default values for priors, constants, and other model information are obtained using built-in helper functions (Table 2). We provide the philosophy behind our default prior choices in the discussion, but we have found that specifying appropriate custom priors for a given context to be more successful than using the default options.

**Table 2.**
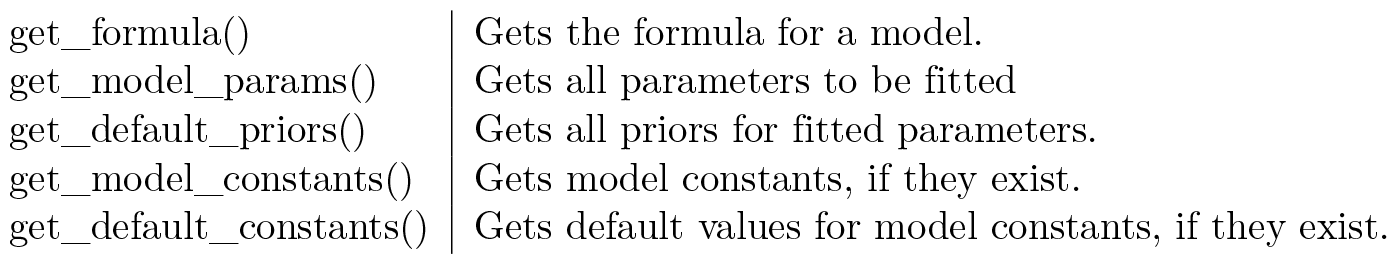
bayesTPC functions to access information on implemented model components.

|  |  |
| --- | --- |
| `get_formula()` | Gets the formula for a model. |
| `get_model_params()` | Gets all parameters to be fitted |
| `get_default_priors()` | Gets all priors for fitted parameters. |
| `get_model_constants()` | Gets model constants, if they exist. |
| `get_default_constants()` | Gets default values for model constants, if they exist. |

In addition to changing prior specification, other MCMC control parameters can be adjusted as arguments to `b_TPC()`, including the number of iterations (`niter`) or burn-in period length (`burn`). Also, any of four sampling methods implemented in `nimble` can be specified using the `samplerType` argument.

### 4.4 MCMC Diagnostics

`bayesTPC` provides MCMC diagnostic plots to support a variety of posterior checks. Chain convergence needs to be confirmed for analyses that depend on MCMC samples to be reliable. The `traceplot()` function wraps `coda::traceplot()` and shows the sampled values by sequential iteration. If `burn` has been specified, `traceplot()` only shows the samples after the burn-in period.

Another important task in Bayesian analysis is to examine the relationship between the prior information and the posterior samples. The `ppo_plot()` function shows the degree of overlap between the priors specified versus a kernel density estimation of the posterior sample. This enables users to determine whether data are informative for a particular parameter, and some indication of how strongly the prior information has influenced the analysis.

### 4.5 Summaries and visualization

The fitted object is returned by `b_TPC()` provides information about the fitted model (including data, model formula, and priors) with the samples from the posterior distribution. Generic methods are provided for `print()` and `summary()`. `print()` provides a quick overview of the fitted model while `summary()` gives a detailed summary of the MCMC results and returns marginal center (mean or median) and bounding (95% quantiles or HPD intervals) based on the MCMC samples. We additionally provide the maximum *a posteriori* (MAP) estimator – the MCMC sample with the highest posterior probability among the obtained samples, similar to a classical maximum likelihood estimator.

It is also possible to plot the marginal posterior and joint distributions. As noted above, marginal priors can be plotted with the marginal posterior using `ppo_plot()`. The `bayesTPC_ipairs()` function wraps `IDPmisc::ipairs()` to visualize the (pairwise) joint posterior distribution of parameters using default colors to represent the relative density of samples.

We also provide visualizations of the fitted TPC’s posterior distribution. The defaults for the (generic method) `plot()` function plots the median and 95% Highest Posterior Density (HPD) interval of the *fitted function* (i.e., plugging the samples into the TPC function, and calculating the median and HPD interval at all evaluated temperatures). Alternatively one may use `posterior_predictive()` with `plot_prediction()`. The `posterior_predictive()` function simulates draws from the posterior predictive distribution, so it includes both the samples describing the TPC function and the observational model. It then uses these samples to calculate the mean/median and the HPD interval of those simulated points. The output of `posterior_predictive()` can be fed into `plot_prediction()` to visualize the these posterior predictive distribution.

## 5 Example: Longevity data for *Aedes aegypti* mosquitoes

### 5.1 Data

To demonstrate the basic workflow of `bayesTPC`, we fit TPCs to individual-level data on adult longevity in *Aedes aegypti* [5]. These data can be loaded into `R` from the VecTraits database (https://vectorbyte.crc.nd.edu/vectraits-explorer) using `get_datasets()` a helper function that interacts with VecTraits using dataset id numbers for retrieval.

As described in Section 4.3, trait data must be stored as a `list` with names `Trait` for the response and `Temp` for the corresponding temperatures (in ^*°*^C) before fitting is performed:

### 5.2 Inference with default settings

Once the data are formatted, we can fit a TPC to the dataset using default settings with a single function call. Adult longevity are numeric data where a concave down unimodal response, such as a Briere function, is likely appropriate.

~~~
bayesTPC Model Specification of Type:
  briere
Model Formula:
  m[i] <- (q * Temp * (Temp - T_min) * sqrt((T_max > Temp)* abs(T_max - Temp))
* (T_max > Temp) * (Temp > T_min))
Model Distribution:
Trait[i] ∼ T(dnorm(mean = m[i], tau = 1/sigma.sq), 0,)
Model Parameters and Priors:
  q ∼ dunif(0, 1)
  T_max ∼ dunif(25, 60)
  T_min ∼ dunif(0, 24)
Prior for Variance:
  sigma.sq ∼ dexp(1)
~~~

We fit the Briere model using the `b_TPC` function with arguments corresponding to the data object and the name of the TPC model that we want to fit.

Once the fitting process completes, we can examine the fitted model object using `print`. This provides details about the model fit, priors, and simple summaries of the MCMC samples.

~~~
bayesTPC MCMC of Type:
  briere
Formula:
  m[i] <- (q * Temp * (Temp - T_min) * sqrt((T_max > Temp) * abs(T_max - Temp))
* (T_max > Temp) * (Temp > T_min))
Distribution:
Trait[i] ∼ T(dnorm(mean = m[i], tau = 1/sigma.sq), 0,)
Parameters:
                  MAP    Mean Median      Priors
T_max          34.464  34.453 34.410 dunif(25, 60)
T_min           0.000   2.559  1.730 dunif(0, 24)
q               0.005   0.006  0.005 dunif(0, 1)
sigma.sq       16.643  17.034 17.001   dexp(1)
~~~

### 5.3 Plots

`b_TPC()` returns an object of class `btpc_MCMC` containing model specification information, data, and the MCMC samples (as an `mcmc` object from the package `coda` [17]). Before using or interpreting a fitted model, users should check the MCMC traceplots to ensure chains have converged. If the model has converged, the marginal traceplot will eventually start varying around a single value, resembling a “fuzzy caterpillar”.

**Figure 1.**
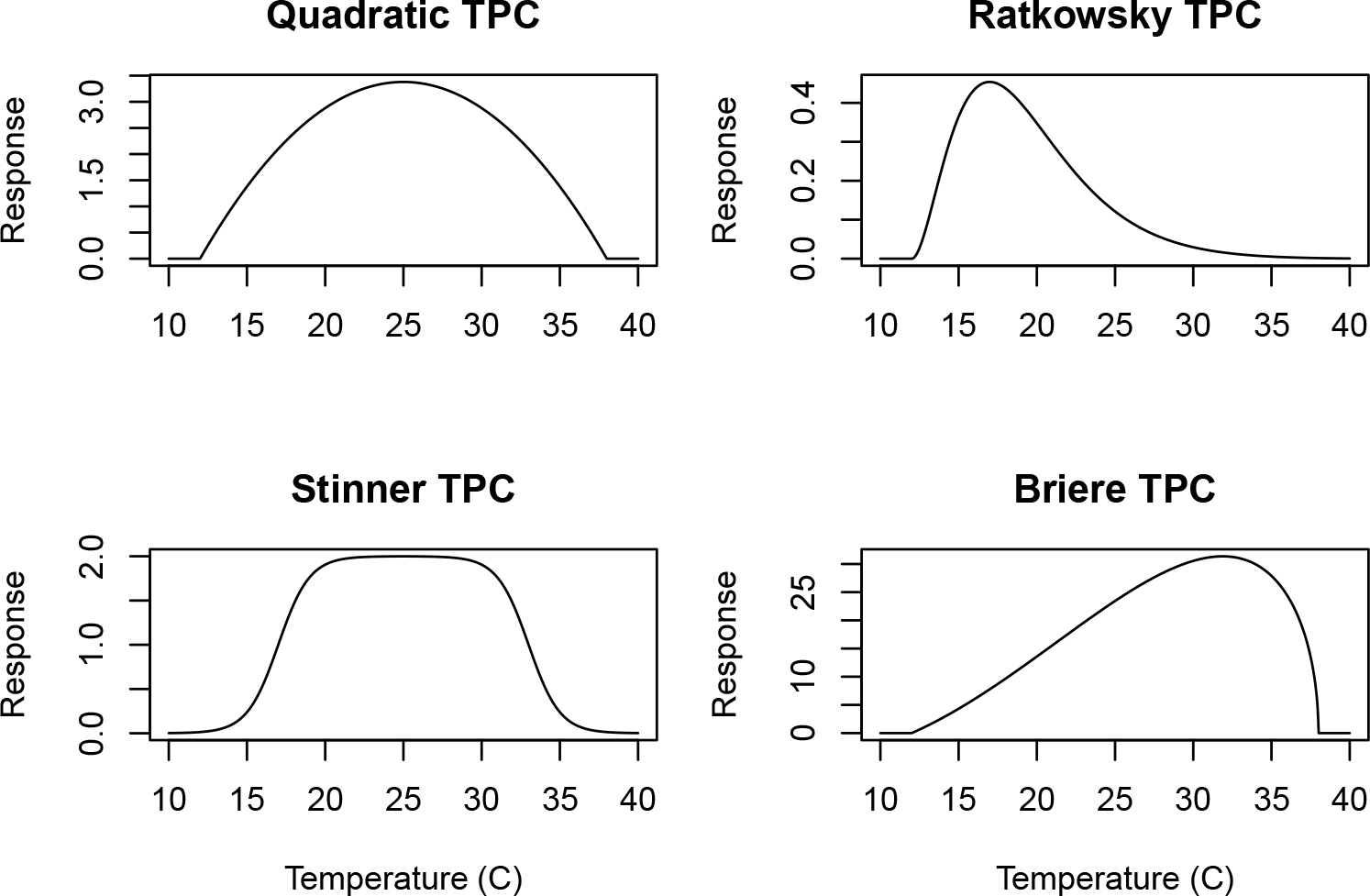
Examples of some TPC model shapes implemented in `bayesTPC`.

The chains in Fig 2 are slow to converge, so we specify a burn-in period and only consider samples obtained after burn-in. Here, a value of 5000 should be sufficient. We can re-visualize with this burn-in (by adding burn=5000 as an argument to traceplot):

**Figure 2.**
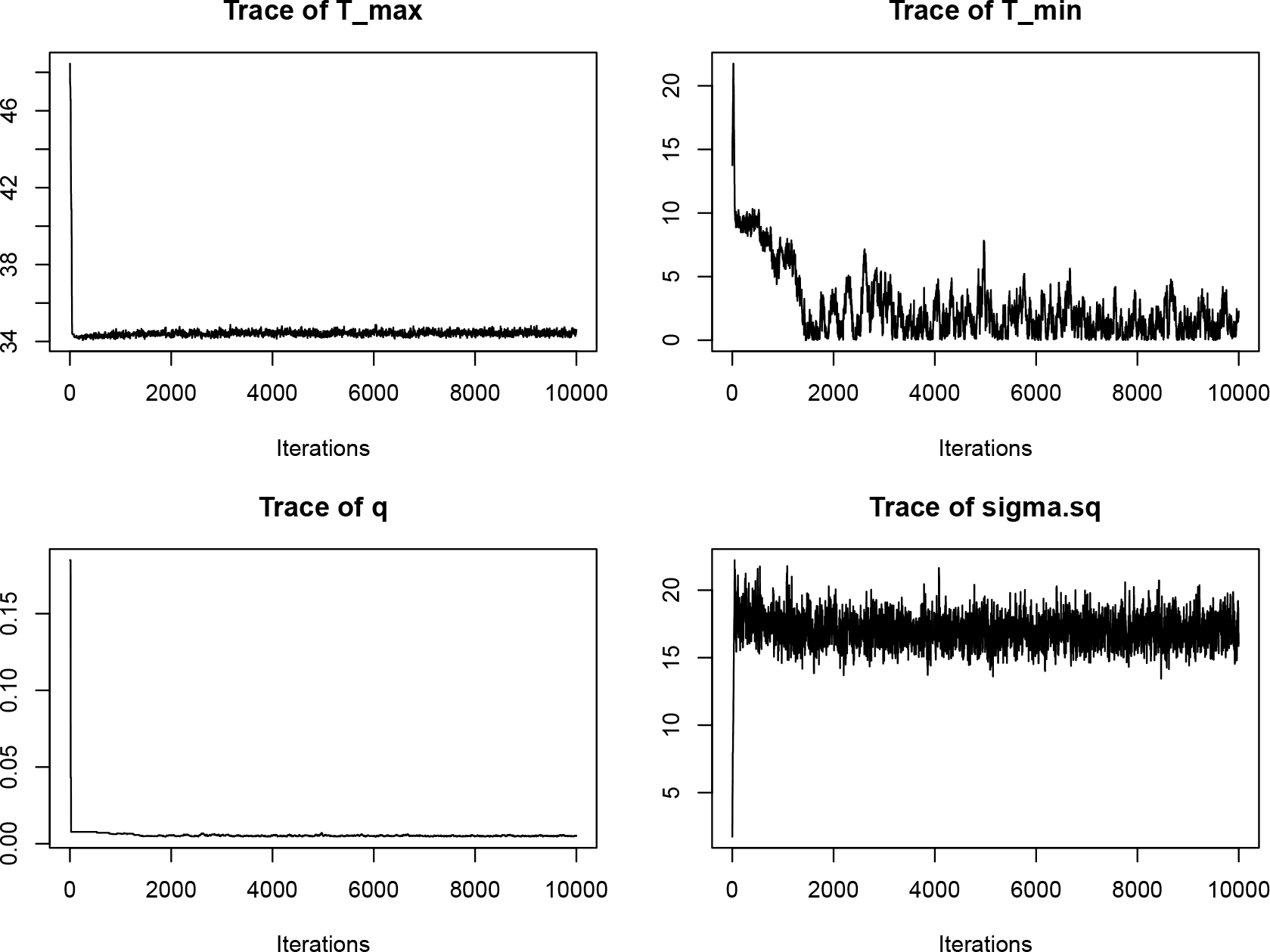
Traceplots for the three parameters of the Briere TPC and the observation parameter for model fitted to the adult longevity data.

**Figure 3.**
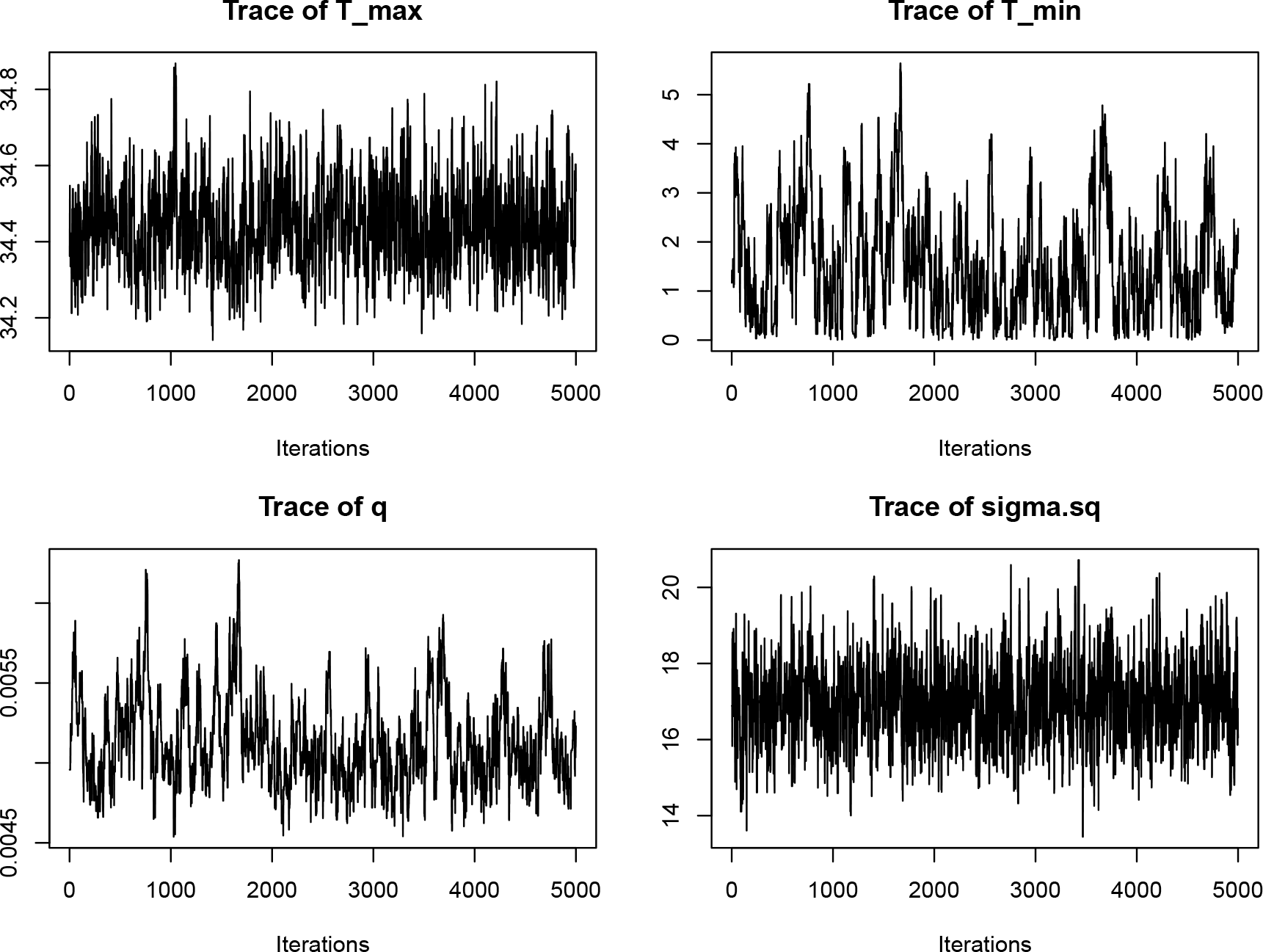
Traceplots for the Briere model parameters and the observation parameter for model fitted to the adult longevity data. Here, the burn-in portion of the MCMC chains has been dropped.

All of the chains now have the desired “fuzzy caterpillar” look. At this point, we would usually re-run the original fitting function, specify the burn-in and increase the total sample size to ensure sufficient samples. This approach drops the burn-in samples from the returned `btpc_MCMC` object.

With appropriate burn-in values selected, we can use `plot()`, and `posterior_predictive()` with `plot_prediction()` to examine the fit of the model in two ways (Fig 4).

**Figure 4.**
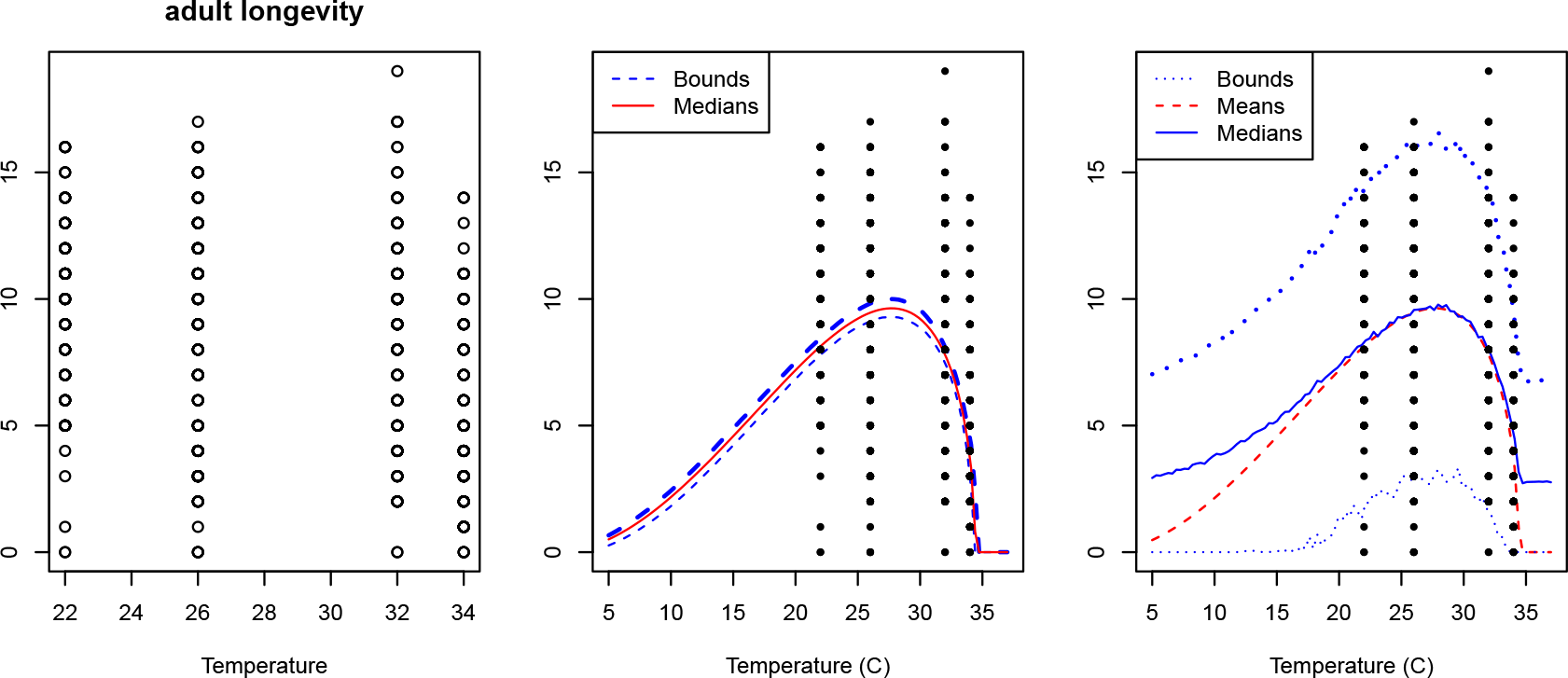
Plots of the adult longevity data (LEFT) with a comparison of the plots produced by plot() (CENTER) and the combination of `posterior_predictive()` and `plot_prediction()` (RIGHT). By default, these only make plots/predictions within the range of temperatures included in the fitted data. The `temp_interval` argument can be used to plot predictions across a broader temperature range.

**Figure 5.**
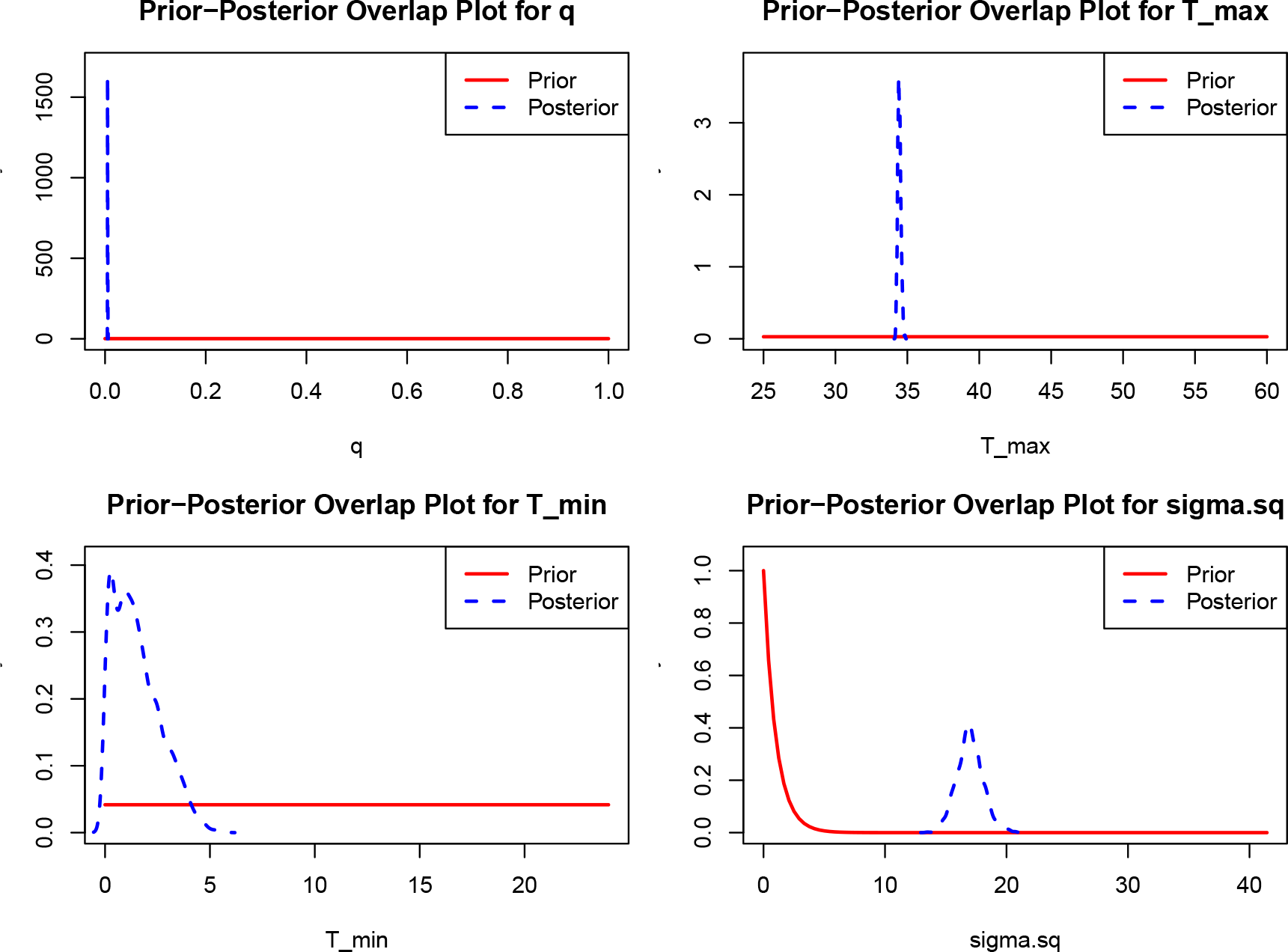
Marginal prior/posterior for the Briere TPC parameters and the observation parameter for the model fitted to the adult longevity data. The burn-in was dropped using the `burn` argument.

**Figure 6.**
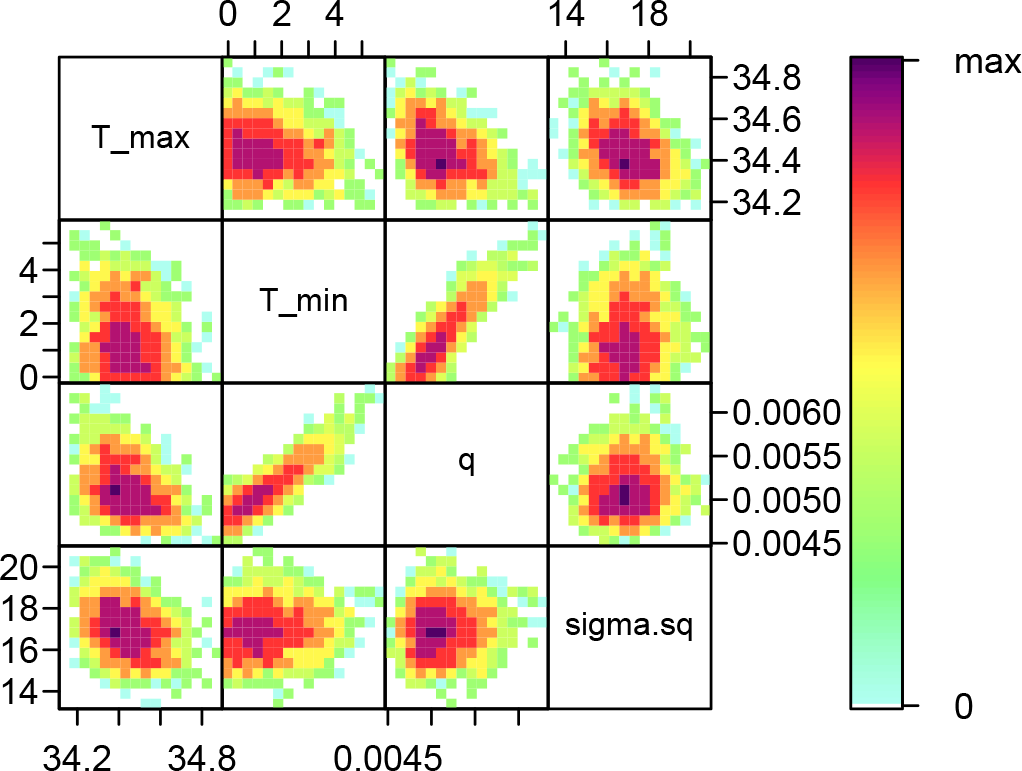
Pairwise visualization of the joint posterior distribution of parameters for the Briere TPC parameters and the observation parameter for the model fitted to the adult longevity data. The burn-in is dropped for this plot using the `burn` argument.

We can obtain marginal priors and posteriors of our model parameters with `ppo_plot()`. This enables us to confirm that our posterior distributions have been informed by the data.

Finally, we can visualize the joint posterior distribution of parameters to understand how estimates are related to each other using `bayesTPC_ipairs()`.

### 5.4 Summaries

If we are satisfied with the traceplots, fits, and prior/posterior plots, users may want to examine additional summary output (for example to make tables) or to save summaries. Numerical summaries are available through the `print()` function shown above, but we can see more details using `summary()`.

~~~
bayesTPC MCMC of Type:
 briere
Formula:
 m[i] <- (q * Temp * (Temp - T_min) * sqrt((T_max > Temp) * abs(T_max - Temp))
* (T_max > Temp) * (Temp > T_min))
Distribution:
Trait[i] ∼ T(dnorm(mean = m[i], tau = 1/sigma.sq), 0,)
Priors:
 q ∼ dunif(0, 1)
 T_max ∼ dunif(25, 60)
 T_min ∼ dunif(0, 24)
 sigma.sq ∼ dexp(1)
Max. A Post. Parameters:
     T_max   T_min       q  sigma.sq   log_prob
   34.4643  0.0005  0.0048   16.6435 -1925.7825
MCMC Results:
Iterations = 1:10000
Thinning interval = 1
Number of chains = 1
Sample size per chain = 10000
1. Empirical mean and standard deviation for each variable,
  plus standard error of the mean:
               Mean       SD   Naive SE Time-series SE
T_max     34.452539 0.609009 6.090e-03        0.0357657
T_min      2.558671 2.620893 2.621e-02        0.4400780
q          0.005706 0.006834 6.834e-05        0.0002871
sigma.sq  17.034134  1.218265 1.218e-02       0.0379541
2. Quantiles for each variable:
                2.5%              25%              50%              75%              97.5%
T_max      34.197591        34.337738        34.409928        34.496010         34.671707
T_min      0.069499          0.768908         1.729905         3.317182          9.403816
q          0.004698          0.004962         0.005172         0.005556          0.007711
sigma.sq  14.951127         16.302953        17.000513        17.728418         19.439654
~~~

We may want to store some of the sample statistics for later use. The sample-based MAP estimator is calculated and saved as part of our fitting process, and can be obtained directly from the fit object.

~~~
   T_max       T_min          q   sigma.sq    log_prob
34.46433     0.00048    0.00480   16.64349 -1925.78250
~~~

### 5.5 Comparing and Selecting Models

Often we are uncertain of what is the preferred shape for a TPC (e.g., left- or right-skewed, symmetric, etc). In these cases we may want to fit (and check) multiple model fits and then choose between them. For the example data we fit two additional functional forms, quadratic and Stinner (Fig 7).

**Figure 7.**
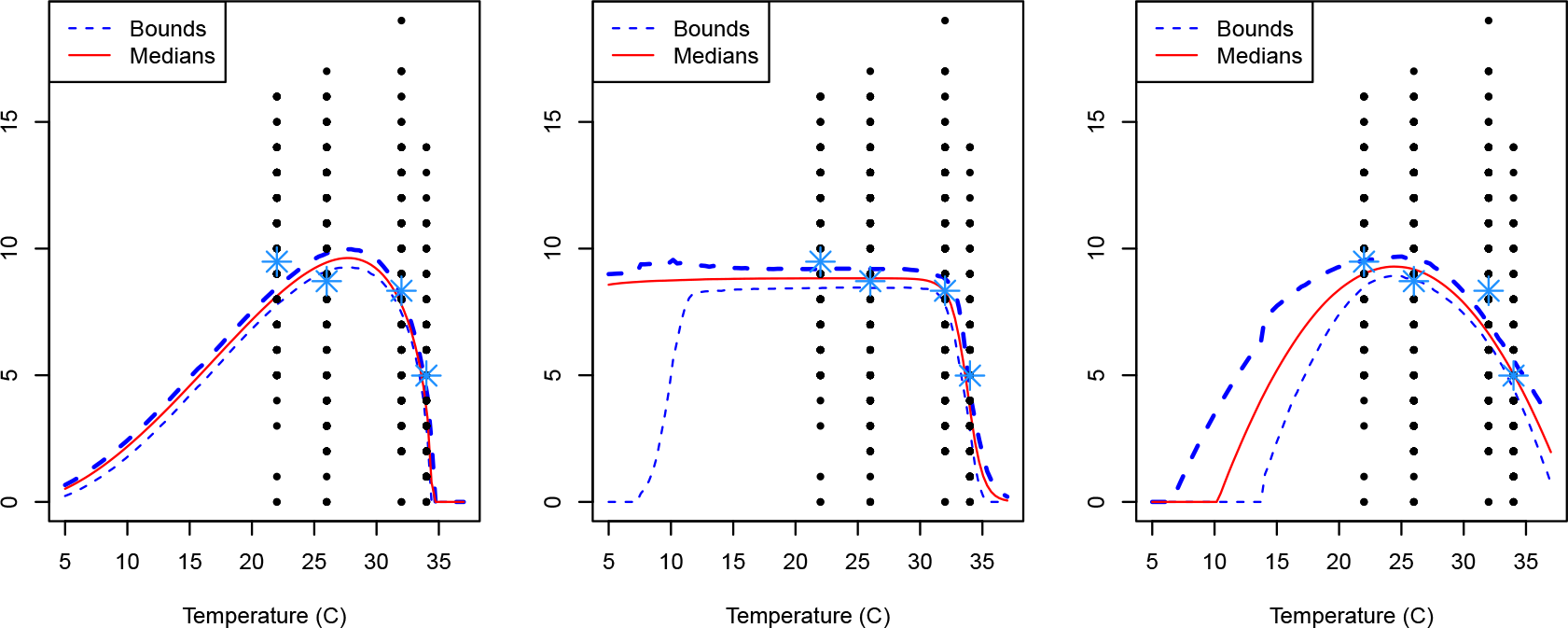
Observed adult longevity data with plots produced by the plot() function for the Briere (left), Stinner (center), and quadratic (right) model fits. Blue stars denote mean trait values at each temperature.

Although, *a priori* we would expect the quadratic fit to be best (previous work suggests it is a good choice for lifespans), all of the fits seem visually reasonable. wAIC values can be obtained with `get_WAIC()` and used compare our fitted models. The preferred model will be the one with the lowest wAIC value.

~~~
  WAIC      lppd   pWAIC
3811.769384 -1902.876741 3.007951
  WAIC      lppd   pWAIC
3825.437397 -1909.197548 3.521151
  WAIC      lppd   p WAIC
3793.919864 -1893.008227 3.951705
~~~

The output here indicates that the Stinner model is the best/most parsimonious fit, even though it has more parameters. However, more data at lower temperatures are likely needed to more convincingly choose between the three models.

## 6 Discussion/Conclusion

We developed `bayesTPC` to support the adoption of Bayesian approaches in thermal biology research, and promote TPC fitting that adequately quantifies uncertainty. To make Bayesian TPCs easy to fit, we implemented sets of default TPCs, likelihoods, and priors that are well-suited for non-negative performance data that consist of averages across individuals/replicates. The methods implemented here can also be used for individual-level data, and for count (Poisson) or success/failure data (Binomial or Bernoulli). For more advanced practitioners, we also provide approaches and guidance for extensibility of models and likelihoods.

We set reasonable defaults to allow straightforward fitting, but we acknowledge that our choices could be overly restrictive for some users. For example, many TPC models include *T*_max_ and *T*_min_ parameters that determine where the TPC reaches zero. These functional forms require that *T*_max_ *> T*_min_. We therefore set non-overlapping uniform priors for these parameters to ensure the inequality is preserved for any sampler. This is a convenient and robust choice for this purpose. We suggest that most users simply change the bounds to modify these priors. However, because support for the uniform is finite, the posterior probability of any settings outside the prior is zero. Thus, the sampler cannot reach a “true” value of a parameter if it is outside the chosen range. Additional inspection is required to decide whether changes to particular prior bounds are needed.

Uniform priors are also highly restrictive in terms of the shape that we assume for *T*_max_ and *T*_min_. These priors assume that any value within the given range is equally likely. If, at the outset, we have little information, this is not a terrible assumption. However, if prior information about these limits is available, we may want to assume a different shape. In this case, users can specify different priors for these parameters but they should be chosen carefully to prevent the sampler violating the condition that *T*_max_ *> T*_min_ (as most other commonly used priors have infinite support).

Finally, we include two additional features in `bayesTPC`. First, the MAP estimator can be obtained as a model summary. We maintain that the full posterior is superior for most purposes. However, the MAP estimator is likely to be the best summary for users requiring single point estimates for use in follow-on modeling exercises (e.g., as an input into a differential equation model). This is because posterior distributions may not be symmetric, and in some cases marginal means of parameters may lie away from the bulk of the posterior distribution, and thus be poor representations of the behavior of the fitted TPC. Second, `bayesTPC` users can obtain wAIC values for fitted models. This is in contrast to other pipelines for Bayesian TPC fitting that rely on deviance information criterion (DIC) for model selection. We choose wAIC as recent work suggests that DIC is less theoretically grounded and more likely to have stability issues than wAIC [18].

## 7 Acknowledgements

LRJ, JWS, and PJH were funded on NSF DMS/DEB #1750113. LRJ, PJH, and SS were funded on NSF DBI #2016264. We thank the participants of the 2023 VectorByte Training Workshop for very helpful feedback on earlier versions of the package.

## 8 Author Contributions

SS: Methodology, Software, Data Curation, Visualization, Validation, Writing - Original Draft, Writing - Review & Editing; JWS: Methodology, Software, Writing - Review & Editing; PJH: Data Curation, Validation, Visualization, Writing - Review & Editing; LRJ: Conceptualization, Methodology, Visualization, Validation, Writing – Review & Editing

